# Assortative mating can impede or facilitate fixation of underdominant alleles

**DOI:** 10.1101/042192

**Authors:** Mitchell G Newberry, David M McCandlish, Joshua B Plotkin

## Abstract

Although underdominant mutations have undoubtedly fixed between divergent species, classical models of population genetics suggest underdominant alleles should be purged quickly, except in small or subdivided populations. Here we study the fixation of underdominant alleles at loci that also influence mate choice, such as loci encoding coloration patterns visible to mates and predators alike. We analyze a mechanistic model of positive assortative mating in which individuals have *n* chances to sample compatible mates. This one-parameter model naturally spans the two classical extremes of random mating (*n* = 1) and complete assortment (*n* → ∞), and yet it produces a complex form of sexual selection that depends non-monotonically on the number of mating opportunities, *n*. The resulting interaction between viability selection and sexual selection can either inhibit or facilitate fixation of underdominant alleles, compared to random mating. As the number of mating opportunities increases, underdominant alleles can fix at rates that even approach the neutral substitution rate. This result is counterintuitive because sexual selection and underdominance each suppress rare alleles in this model, and yet in combination they can promote the fixation of rare alleles. This phenomenon constitutes a new mechanism for the fixation of underdominant alleles in large populations, and it illustrates how incorporating life history characteristics can alter the predictions of population-genetic models for evolutionary change.

## 1. INTRODUCTION

An allele is underdominant, or overdominant, if it experiences reduced, or enhanced, fitness as a heterozygote compared to either homozygote. Overdominance has been intensively studied as a mechanism of maintaining diversity in populations, whereas underdominance reduces diversity and has been studied as a mechanism for population differentiation and speciation[Wright, 1941; Lande, 1979].

Underdominance typically occurs when the two homologous gene copies at a diploid locus must act in concert with each other. One classic example is a chromosomal alteration that disrupts meiosis in heterozygotes [Lande, 1979]. Although much research on the fate of underdominant alleles has focused on chromosomal rearrangements, underdominant alleles occur and have evolutionary consequences in many other contexts. Underdominance has been observed at loci that regulate gene expression[Smith *et al.*, 2011; Stewart *et al.*, 2013], and engineered under-dominant transgenes play an important role in strategies to control insect disease vectors[Curtis, 1968; Sinkins and Gould, 2006; Reeves *et al.*, 2014]. Underdominance has also been observed at loci controlling quantitative traits, such as body size[Kenney-Hunt *et al.*, 2006], that are known to influence mate choice [Crespi, 1989]. Likewise, heterozygote deficits in hybrid zones have been observed at the loci encoding color patterning in *Heliconius* butterfly species, where coloration has been implicated both in assortative mating [Jiggins *et al.*, 1996; Arias *et al.*, 2008] and also in viability via the avoidance of predators [Mallet and Barton, 1989; Kapan, 2001; Langham and Benkman, 2004]. Motivated by these examples, we focus here on the fate of alleles that simultaneously influence mate choice and viability. We ask whether assortative mating will facilitate or impede the fixation of an underdominant allele.

In general, the fixation of an underdominant allele is exceedingly rare, at least in theory[Wright, 1941; Kimura, 1962; Ortíz-Barrientos *et al.*, 2007]. A classical approximation due to Lande [1979] for the probability of fixation *u* of a novel underdominant allele with heterozygote disadvantage *s* in a well-mixed population of size *N* is:

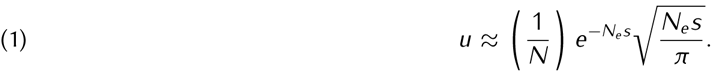

This fixation probability decreases exponentially with the effective population size, *N*_*e*_. Under this analysis fixation through drift of a novel underdominant allele is possible only when the effective population size is extremely small — on the order of tens or hundreds of individuals. Nonetheless, empirical observations provide strong evidence that underdominant alleles have indeed fixed in populations.

Several possible resolutions to this paradox have been proposed. One common solution is based on Wright’s shifting balance theory[Wright, 1931, 1941, 1982]. According to Eq. 1, underdominant alleles may fix in extremely small populations, so that fixation across a species as a whole might occur through successive fixation in small, mostly-isolated subpopulations[Wright, 1941; Lande, 1979; Slatkin, 1981; Barton and Rouhani, 1991; Whitlock, 2003; Altrock *et al.*, 2011]. However, for this process to occur effectively it is typically necessary to include extinction and recolonization of demes[Lande, 1979; Michalakis and Olivieri, 1993; Roze and Rousset, 2003]. Other theoretically possible explanations[Hedrick, 1981] include meiotic drive and partial selfing [Charlesworth, 1992]. Other authors have investigated linkage to locally adaptive alleles[Navarro and Barton, 2003; Kirkpatrick and Barton, 2006].

Here we consider mate choice as a mechanism to explain the fixation of underdominant alleles. This mechanism is limited to loci that simultaneously influence mate choice and also viability. Phenotypes that serve as mating cues are often also subject to surveillance by predators, where rarity is typically detrimental to survivorship. In the context of coloration patterns in the *Heliconia*, for example, rare phenotypes often both experience increased predation and also determine assortment [Mallet and Barton, 1989; Kapan, 2001; Langham and Benkman, 2004; Jiggins *et al.*, 1996; Arias *et al.*, 2008]. Likewise, in vertebrates ranging from cichlids [Sefc *et al.*, 2014; Anderson *et al.*, 2015] to finches [Blount, 2004; Blount *et al.*, 2003], carotenoid coloration phenotypes are well known to influence mate choice and viability alike. Therefore, although this study does not apply to chromosomal rearrangements, which are unlikely to induce assortment, it does apply to a broad range of other loci and underdominant mutations.

A broad literature has successfully addressed questions about diversity of mating systems in nature[Andersson, 1994; Andersson and Simmons, 2006; Clutton-Brock and McAuliffe, 2009], their evolutionary maintenance and optimality[Lande, 1981; Lande and Schemske, 1985; Real, 1990; Goodwillie *et al.*, 2005; Kokko *et al.*, 2006; Jones and Ratterman, 2009; Wiegmann *et al.*, 2010], and their consequence for allele frequency change [e.g. inbreeding depression Charlesworth, 1992; Nagylaki, 1992; Whitlock, 2000]. While there are many analytical studies on the effects of mating systems on allele dynamics, they tend to provide either a deterministic treatment under a specific model of mate choice[Karlin, 1978; O’Donald, 1980; Kirkpatrick, 1982; Seger, 1985; Otto *et al.*, 2008], or a full stochastic treatment but only for mating systems that are essentially equivalent to a fixed population structure [i.e. a constant inbreeding coefficient, *F*, Caballero and Hill, 1992; Damgaard, 2000; Roze and Rousset, 2003; Glémin, 2012].

Rather than stipulate a fixed population structure or a constant probability of selfing, we will provide a stochastic analysis of a mechanistic model of assortative mating. The model is defined defined in terms of the absolute number of individuals, *n*, that an organism can survey before eventually choosing a mate. This one-parameter model of positive assortative mating coincides with classical partial self-fertilization in two limiting cases. For *n* = 1 the model corresponds to random mating, whereas as *n* → ∞ it corresponds to complete assortment. Although the fixation probability of a new mutation under partial selfing smoothly interpolates between these two extreme cases[Charlesworth, 1992], we will show that the fixation probability in our model has a non-monotonic dependence on the life-history parameter, *n*. Increasing the number of mating opportunities beyond *n* = 1 initially inhibits the fixation of underdominant alleles; but increasing *n* yet further eventually facilitates fixation, allowing rates approaching that of neutral substitutions. These results are surprising because the mate choice model induces a form of positive frequency-dependent sexual selection that, in the absence of viability selection, always inhibits the fixation of rare alleles. We will explain these results in terms of the geometry of a slow manifold that arises under preferential mating, analogous to the Hardy-Weinberg equilibrium for random mating, and we discuss implications for the evolution of underdominant alleles in nature.

## 2. A MECHANISTIC MODEL OF MATE CHOICE

Models of mate choice, assortative mating, and sexual selection have been extensively studied and characterized[Gavrilets, 2004]. In partial self-fertilization or mixed mating models, individuals mate with themselves with a fixed probability and the mating system does not alter allele frequencies[Haldane, 1924]. In partial assortative or preferential mating models, parents prefer to mate with their own genotype, or with particular other genotypes, and the mating system itself can alter allele frequencies by sexual selection. Such models are typically formulated[Karlin, 1978] by specifying, exogenously, the chance that one genotype will mate with another genotype, which allows mating with rare types according to preference regardless of the frequency of the rare type. This formulation implies that individuals are able to census all other individuals in the entire population in the decision to mate — which is unrealistic for many biological populations. Even some models that incorporate a search cost[Otto *et al.*, 2008] still have the property that the cost of finding vanishingly rare types is fixed, regardless of their frequency.

Here we study a one-locus, two-allele model of hermaphroditic diploid individuals in which parents prefer to mate with their own genotype. Over the course of successive discrete generations we track the frequencies of all three diploid genotypes, *x*_*i*_ for *i* ∈ {*aa, aA, AA*}, in a population of constant size *N*. A parent can mate with any individual from a pool of *n* prospective mates, drawn uniformly with replacement from the population. If there is a mate of the parent’s own genotype among these *n* prospective mates, then the parent chooses that mate; otherwise, the parent chooses uniformly at random from the pool of *n* prospective mates. The mating always produces one offspring. This model is equivalently described as parents having *n* chances to find a compatible mate by sampling randomly from the population. Parents sample mates from the population up to *n* times, mating immediately with any individual of their own genotype, or ultimately accepting any genotype on their *n*th chance. Because all parents choose among *n* potential partners, we call this the *n*-choice model. Several similar models, proposed by O’Donald [1980] and Janetos [1980], have previously been analyzed in a deterministic setting [O’Donald, 1980; Seger, 1985].

The *n*-choice model is a simple, mechanistic implementation of positive assortative mating that accounts for the fact that in reality rare genotypes are less likely to find preferred mates. Importantly, the model does not rely on an intrinsic capacity for selfing. In other words, there is no fixed chance that an individual will reproduce with its own genotype regardless of the frequency of that genotype. Rather, individuals census a limited number of possible mates from the population with replacement. While this sampling scheme includes a small probability that an individual will ‘encounter’ and mate with itself, we demonstrate robustness of the results to this and other biological assumptions in the Discussion section.

The outcomes of mating are dependent on all genotype frequencies, and so we must tabulate the probability of each mating[Nagylaki, 1992]. The *n*-choice model does not explicitly distinguish between sexes, as with hermaphroditic or monoecious populations. Nevertheless, we may consider the mate-choosing parent in any pairing as the “female” or macrogamete-donor parent. According to the model, the probability that a female (i.e. mate-choosing) parent with diploid genotype *i* finds a mate of her own genotype is (1 − (1 − *x*_*i*_)^*n*^). Otherwise, the parent reproduces with a different genotype with probability proportional to that genotype’s frequency in the population. The probability distribution of mating types *G* conditional on the female’s genotype *P* is thus

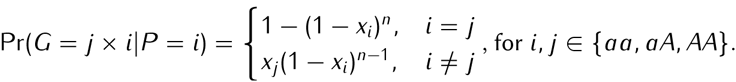

The genotypic distribution of offspring from a given mating pair is Mendelian. We can therefore compute the distribution of zygotic genotypes *Z* after mate choice and reproduction by conditioning on the distribution of matings:

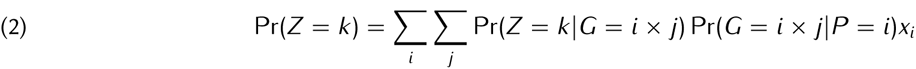

where Pr(*Z* = *k|G* = *i* x *j*) denote the standard Mendelian probabilities, and *i, j, k* range over the three diploid genotypes *aa, aA*, and *AA*.

Following mate choice and production of a large zygotic pool we assume that viability selection modifies the frequencies of zygotic genotypes. The subsequent generation of reproductive adults is then drawn from the post-selection zygotic pool. Assuming the zygote pool is very large relative to the population size, the genotype of each surviving sampled adult in the next generation is drawn from the trinomial, fitness-weighted zygote distribution. Genotype *i* is drawn with probability

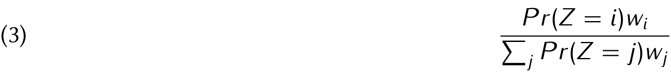

The fitness scheme for underdominant alleles can be expressed as *w*_*aa*_ = *w*_*AA*_ = 1, and *w*_*aA*_ = 1 − *s*.

## 3. Analysis

We explore the influence of mate choice on the fixation rate of alleles by analyzing the *n*-choice assortative mating model in finite populations.

In a finite population of constant size *N* adults, the frequencies of the adult genotypes of the next generation are drawn from the trinomial distribution with the probability of genotype *i* given by Eq. 3. We denote the outcome of this trinomial draw, for each genotype *i* ∈ {*aa, aA, AA*}, by 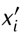. In other words, the values 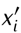 denote the frequencies of the three genotypes in the next generation of adults, given the frequencies *x*_*i*_ in the current generation.

The expected frequency of genotype *i* among adults in the next generation is given simply by the frequency of that genotype in the post-selection zygotes, that is by Eq. 3 above, which we henceforth denote E(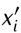). Under the multinomial sampling assumption the variance in the frequency of genotype *i* among the adult individuals in the next generation is simply E(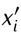) (1 − E(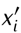))/*N*, and the covariance between different genotypes is — E(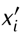) E(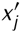/*N*.

Genotype frequencies exist on the simplex *x*_*aa*_ + *x*_*aA*_ + *x*_*AA*_ = 1. We can thus remove one variable from the model by tracking genotype frequencies in some choice of basis for the simplex. Henceforth, as a convenient choice of basis, we will track the frequency of the *a* allele among adults, denoted *p* = *x*_*aa*_ + (1/2)*x*_*aA*_, and one-half the frequency of heterozygous adults, denoted *h* = *x*_*aA*_/2. One-half the frequency of heterozygotes may be thought of as the frequency of the *a* allele present in heterozygotes, and so it has the same units (allele copy frequency) as *p*. The coordinate *h* is preferable to other measures of heterozygosity, such as the quantity *x*_*aA*_/(*p*(1 − *p*)), because it does not depend on allele frequency and it makes no implicit assumption about the shape of the slow manifold.

We can express the expected allele frequency E(*p*′) and the expected heterozygous allele frequency E(*h*′) in the next generation in terms of the current frequencies *p* and *h*. To do so, we first write the mean allele and heterozygote frequencies among zygotes prior to selection as *π* and *η* in terms of *p, q* = 1 − *p* and *h* by expanding Eq. 2:

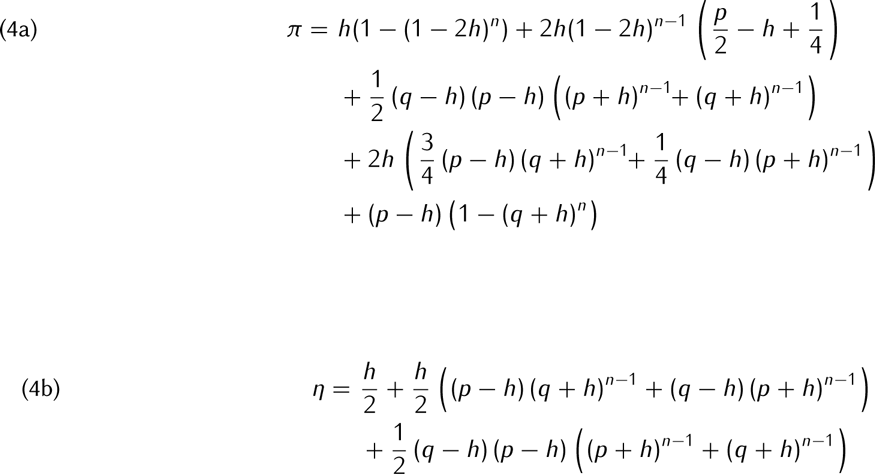

We then use Eq. 3 to obtain expressions for the frequencies of post-selection zygotes in the (*p, h*) basis, which are the expected frequencies of adults in the next generation:

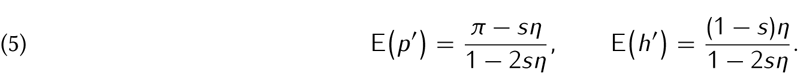

Finally, the variance in adult allele frequencies in the next generation can then be written as

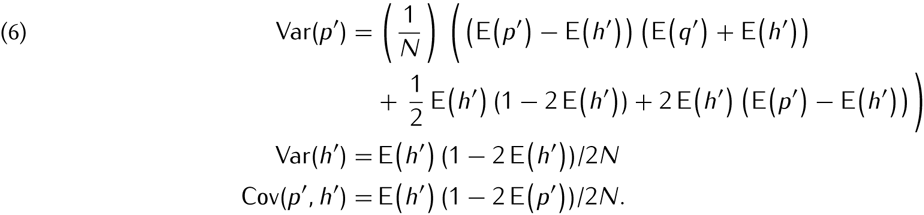

Here we see that Cov(*p*′, *h*′) is positive only if E(*p*′) < 1/2. This expectation and variance will be sufficient to develop a diffusion approximation to the model, along the lines of Kimura[Kimura, 1964].

### 3.1. Behavior in two classical limits

When the sampled pool of prospective mates has only one individual, *n* = 1, or when the pool becomes much larger than the population itself, *n* → ∞, the *n*-choice mating model corresponds precisely to two classical population models: random mating and complete inbreeding, respectively.

When only one prospective mate is allowed per parent, *n* = 1, the model reduces to the classical model of an underdominant allele in a randomly mating population. Eqs. 4 simplify to *π* = *p* and *η* = *p*(1 − *p*). Thus the genotype frequencies among zygotes are at Hardy-Weinberg equilibrium, and viability selection proceeds as in ordinary underdominance with E(*p*′) = (*p* − *sp*(1 − *p*))/(1 − 2*sp*(1 − *p*). When *s* = 0, the dynamics are neutral on 2*N* haplotypes. When there is selection against heterozygotes, *s* > 0, then the fixation probability of a novel allele agrees with Lande’s classic expression, given by equation 1.

On the other hand, when the number of mating opportunities becomes very large, *n* → ∞, preferred genotypes are always available for mating, and the model is equivalent to complete assortment. As we take *n* to infinity in Eq. 4, the frequency of heterozygotes among zygotes approaches *η* = *h/2*. That is, the frequency of heterozygotes is reduced by half at each generation. The population thus rapidly approaches heterozygote frequency zero. Setting *h* = 0 in Eq. 4 and taking *n* to infinity gives *π* = *p*. As selection acts only on heterozygotes, E(*p*′) = *p* and the dynamics are neutral. The variance of *p*′ is E(*p*′) (1 − E(*p*′))/*N*. This variance is the same as that for a population of *N* haplotypes, rather than the 2*N* actually present in the population. Because of complete assortment, each diploid individual behaves roughly as a single haplotype. In this limit of complete assortment, the dynamics of genotype frequencies are always neutral, regardless of *s*, and the fixation probability of a novel allele is always 1/2*N*.

When the number of mating opportunities is intermediate between these two extremes, namely 1 < *n* < ∞, the dynamics of the *n*-choice model are neither neutral nor the same as the dynamics of classical underdominance. Importantly, these dynamics may change allele frequencies dramatically due to a mix of sexual selection and underdominant selection. Analyzing these two-dimensional dynamics requires the development of an appropriate diffusion approximation.

### 3.2. Diffusion approximation

We adopt the techniques used to analyze the fixation probability of an under-dominant allele under random mating to derive a more general expression for the *n*-choice mating model. Under the diffusion limit of Kimura[Kimura, 1964], the probability density *ϕ*(*p, h, τ*) of observing allele frequencies *p* and *h* evolves in time according to the standard Kolmogorov forward equation[Gardiner, 2009], which depends on the instantaneous mean and variance-covariance matrix of the changes in allele frequencies.

We find the instantaneous mean and variance of allele frequency changes, *M*_*i*_ and *V*_*ij*_, by rescaling the discrete model by the population size, *N*. To arrive at a non-trivial diffusion limit we adopt a slight variant of the model above, in which only a fraction *f* of the population undergoes mate choice each generation, while the remainder the population is sampled according to strict clonal reproduction. We take the limit as *N* approaches infinity, scaling *f* and *s* such that *Nf*= *ζ* and *Ns* = *γ* are held constant. The diffusion equation becomes (see *Methods* for derivation)

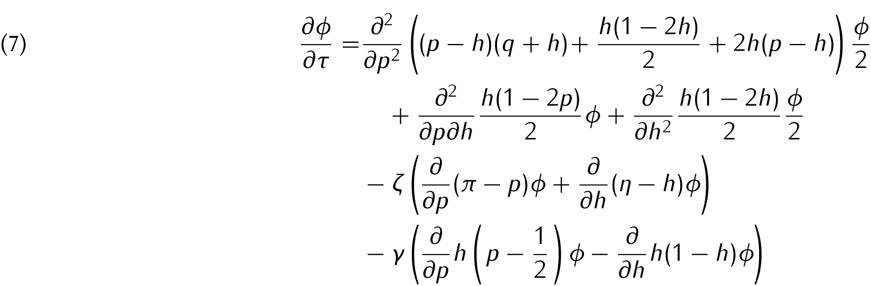

In general, this diffusion in allele-frequency space is two-dimensional and well known to be computationally formidable[Epstein and Mazzeo, 2013]. In the next section we introduce a one-dimensional approximation that makes the diffusion tractable.

### 3.3. Diffusion along the slow manifold

The dynamics in two-dimensional diffusions sometimes approach and remain in the vicinity of a one-dimensional curve, until absorption into a boundary. This behavior can be interpreted as a separation of timescales[Parsons and Rogers, 2015]: there is fast approach to a lower-dimensional manifold, and then slow diffusion along the “slow manifold”. In the case of random mating for a single-locus diploid model, for example, Kimura’s one-dimensional diffusion works because genotype frequencies are assumed to be at Hardy-Weinberg equilibrium at all times. For monoecious random mating, approach to the manifold of Hardy-Weinberg equilibrium takes only a single generation, which is instantaneous in the diffusion timescale. In other mating systems, such as random mating with separate sexes, equilibrium is reached after two generations of mating, or the slow manifold may be approached geometrically. Some dynamics, such as clonal reproduction, do not approach any lower-dimensional sub-manifold whatsoever, and so they exhibit truly two-dimensional diffusions.

Principled approaches to determining the existence and form of the slow manifold are complex [Parsons and Rogers, 2015]. Nonetheless, the dynamics of the *n*-choice mating model clearly exhibit timescale separation in the presence of viability selection against heterozygotes (*s* > 0), participation in the mating system (*f* > 0), or both. The presence of a slow manifold and its width in a finite population can be seen visually in Fig. 1.

**FIGURE 1.**
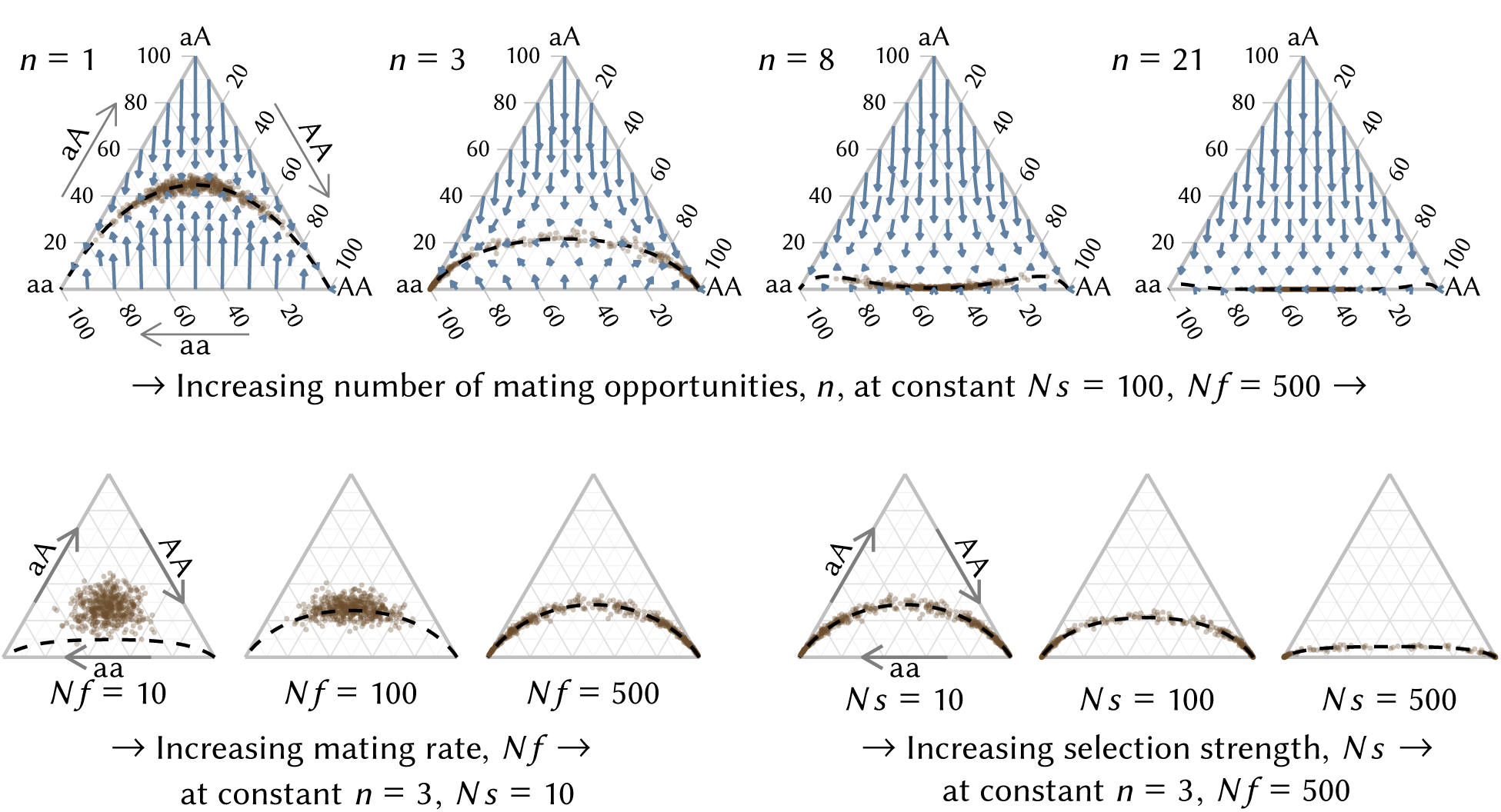
The dynamics of genotype frequencies in the *n*-choice model of assortative mating. Each ternary plot corresponds to a different set of parameter values for the number of mating opportunities, *n*, the per-generation rate of participation in the mating system, *Nf*, and the strength of viability selection against heterozygotes, *Ns*. Blue arrows indicate the expected change in genotype frequencies in one generation. On each plot, tan dots represent the genotype frequencies after 30 generations of stochastic simulation, in 500 replicate populations of size *N* = 1,000 each initialized at the center 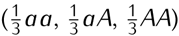. The dynamics quickly converge towards a one-dimensional submanifold within the frequency space. Black dashed lines show the analytically derived position of this one-dimensional manifold, which corresponds to the Hardy-Weinberg equilibrium in the case of random mating (*n* = 1). Increasing the strength of selection, *Ns*, moves the manifold towards zero heterozygosity, while the effect of participation in the mating system, *Nf*, on the shape of the manifold depends on the number of mating opportunities, *n*. In most regimes depicted, information about the initial height (heterozygosity) of the population is lost after 30 generations, as the genotype frequencies have converged to the slow manifold.

We will approximate the two-dimensional diffusion in one dimension by assuming that the dynamics take place along a slow manifold of equilibrium genotype frequencies. This analysis is similar to the classical assumption of convergence to Hardy-Weinberg equilibrium for random mating. However the equilibrium manifold for the *n*-choice model typically entails reduced heterozygosity relative to the Hardy-Weinberg equilibrium (as seen in Fig. 1), either because of strong viability selection against heterozygotes or because assortative matingstend not to produce heterozygotes.

To determine the manifold of equilibrium genotypic frequencies in the *n*-choice model we use the simple principle that frequencies at equilibrium should stay at equilibrium. Mathematically, this implies the condition that the infinitesimal mean change in frequencies (*p, h*) must always be tangent to the slow manifold. Thus, the slow manifold can be defined as a parametric curve (*p*(*l*),*h*(*l*)) such that

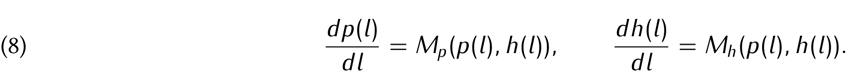

This differential equation has an infinite family of solutions. Fortunately there are additional criteria for equilibrium genotypic frequencies. Since rare alleles are always present as heterozygotes, *dp*(*l*)/*dh*(*l*) must approach *l* as approaches infinity. This criterion in terms of long times is difficult to use in practice, and so we use the symmetry of the dynamics: when *p* = 1/2 the slow manifold is horizontal, meaning E(*h*′) = *h*. We use this symmetry criterion to initialize the parametric curve close to *p* = 1/2. This particular solution to the differential equation above provides a function 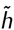(*p*), which defines the manifold of equilibrium heterozygosity as a function of allele frequency.

If we assume that the two-dimensional dynamics in (*p, h*) are restricted to the slow manifold, defined by the curve (*p*, 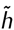(*p*)), then we can treat the dynamics as a one-dimensional diffusion along this manifold satisfying

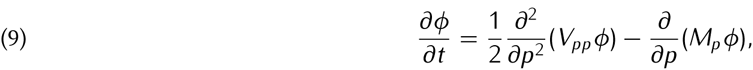

by substituting 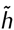(*p*) for *h* in Eq. 4. Although *M*_*p*_ and *V*_*pp*_ depend on *h*, we compute *h* and *η* from *p* by substituting 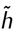(*p*) for *h* in Eq. 4, and we view *M*_*p*_ and *V*_*pp*_ as functions of *p* only. Following Kimura[Kimura, 1962], the solution to a boundary value problem gives the fixation probability *u*(*p*) of a mutant allele initiated at frequency *p* with solution

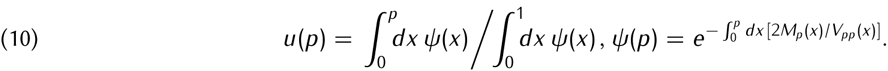

This integral can be computed numerically, giving a good approximation to the fixation probability for arbitrary *n*.

## 4. RESULTS

### 4.1. Sexual selection in the *n*-choice mating model

First we consider the fixation probability of an allele in the absence of viability selection. In the limits of random mating and complete assortment, *n* = 1 and *n* → ∞ respectively, the fixation probabilities are equal to the neutral fixation probability. For an intermediate number of mating opportunities, *n*, however, the mating system itself induces strong sexual selection against rare alleles (cf. Fig. 1), which depresses the fixation probability of novel alleles much below the neutral probability (Fig. 2). And so the resulting fixation probability has a complex, non-monotonic dependence on the number of mates that a parent can survey: increasing *n* beyond one reduces the fixation probability below the neutral value 1/2*N*, but increasing *n* yet further restores the fixation probability until it recovers to the neutral value 1/2*N* in the limit *n* → ∞.

**FIGURE 2.**
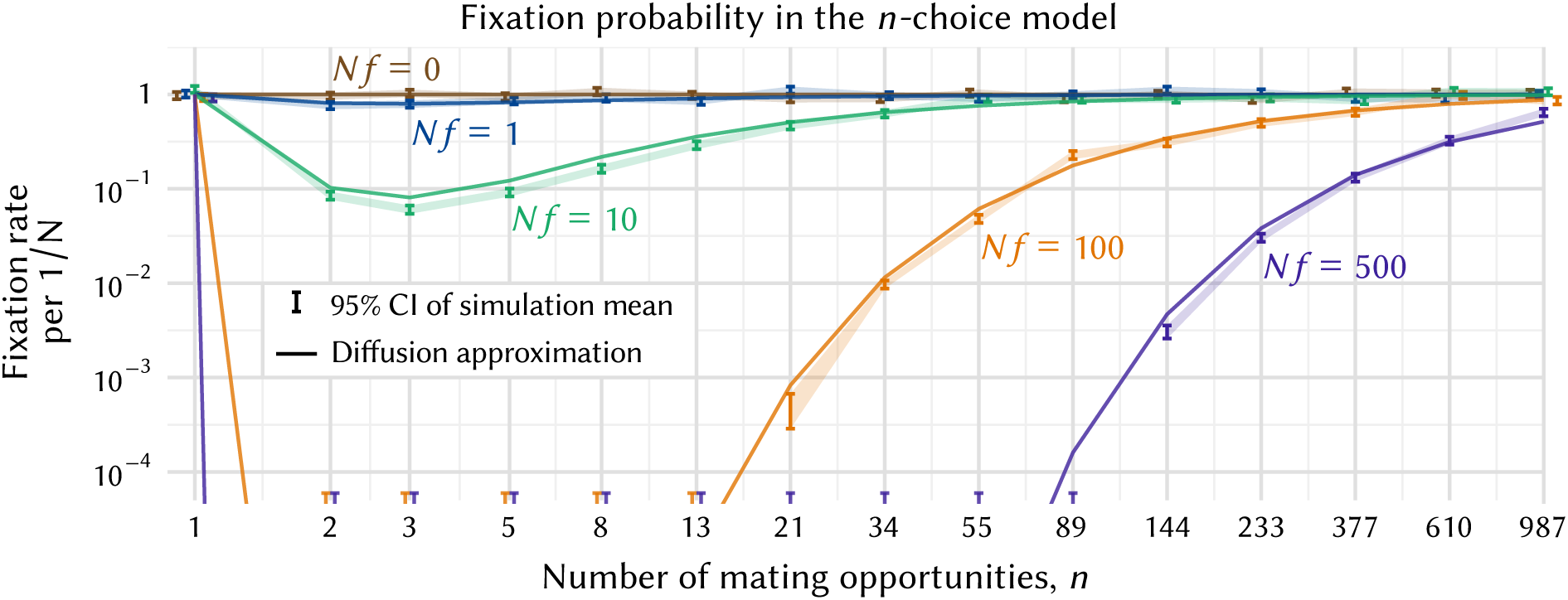
Sexual selection induced by the *n*-choice model of assortative mating. The plot shows the fixation probability of one initial heterozygote in the absence of viability selection, as a function the number of mating opportunities, *n*, and for different rates of participation in the mating system, *Nf*. Solid lines indicate fixation probabilities computed by the diffusion approximation along the slow manifold. Bands and error bars indicate the 95% confidence interval on the mean fixation rate observed in up to 100,000,000 simulated populations of size *N* = 1,000. Overlapping error bars are dodged for clarity. Error bars with no lower bound indicate no fixations observed in 100,000,000 simulations. The fixation probability equals the neutral fixation probability for either *n* = 1 or *n* → ∞. Otherwise, participation in the mating system induces a complex form of sexual selection against rare alleles, whose strength depends non-monotonically on *n*.

The strength of sexual selection against rare alleles depends on both the degree of participation in the mating system, *Nf*, and the number of mating opportunities, *n*. At high mating rates (*Nf* ≥ 100), fixation probabilities at intermediate values of *n* are so low as to be impractical to compute through Monte-Carlo simulation, and they can be known only through numerical integration of the expression derived from the diffusion approximation, Eq. 10. In the regime where Monte-Carlo methods are feasible, the diffusion approximation along the slow manifold is in close correspondence with Monte Carlo simulations across a broad range of values of *Nf*, despite many potential sources of error in the approximation (see Fig. 2).

When the number of potential mates *n* is small but exceeds one, then rare alleles are under-represented among zygotes relative to their parents, as described by Eq. 5. Mendelian inheritance does not alter allele frequencies, and so it is the action of mate choice itself that suppresses rare alleles: parents with common genotypes are likely to find their preferred mates, but parents with rare genotypes are more likely to settle for a common mate. As *n* increases, however, the likelihood of any parent having to settle for a non-preferred mate becomes vanishingly small, and so the differences in mate availability between genotypes become less pronounced. When there is no difference in mate availability even the rarest of parental genotypes can find a mate with their own genotype, and complete assortment ensues.

In summary, Fig. 2 reveals that a mechanistic model of mate choice with two classical limits nonetheless produces a complex form of sexual selection whose strength depends non-monotonically on number of mating opportunities, *n*.

### 4.2. Interaction between sexual selection and underdominant viability selection

What is the fate of novel alleles when we combine the intrinsic effects of *n*-choice mating with viability selection against heterozygotes? Does positive assortative mating facilitate or impede the fixation of an underdominant allele?

The combined effects of viability selection and sexual selection induced by *n*-choice mating are shown in Fig. 3. Here we see that when the number of mating opportunities is small but not one, then preferential mating impedes fixation of an underdominant allele, so that its fixation probability is even lower than the classical prediction of Lande for random mating (*n* = 1). For example, allowing just *n* = 2 opportunities to find a mate with the same genotype reduces the fixation probability of the underdominant allele astronomically.

**FIGURE 3.**
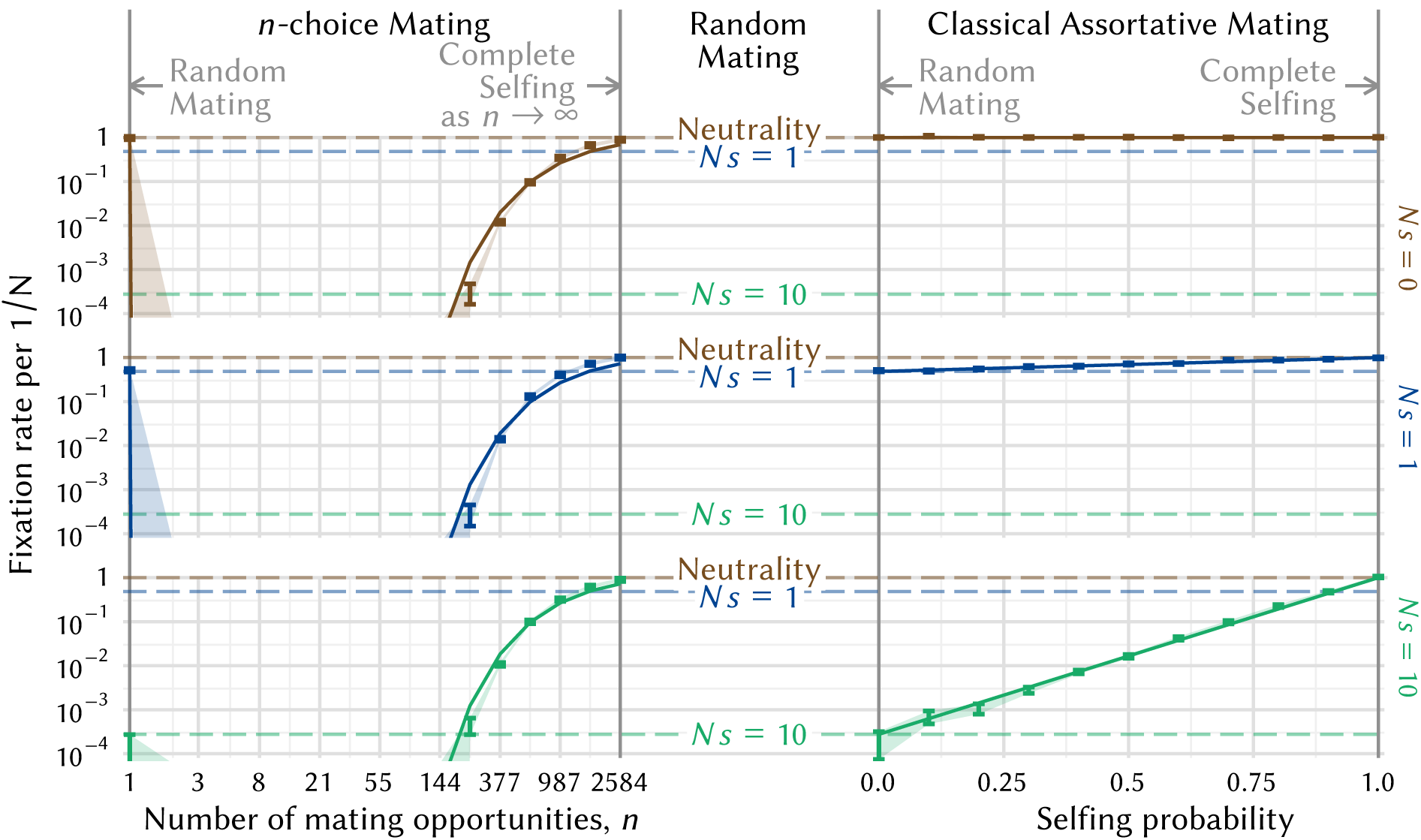
The effect of assortative mating on the fate of an underdominant allele, in the *n*-choice model (left) compared to the classical model of assortative mating by partial self-fertilization (right). The plot shows how the fixation probability of one initial heterozygote depends on the number of mating opportunities, in the case of *n*-choice mating, or on the self-fertilization probability, in classical assortative mating. Vertical bars indicate the 95% confidence interval on the mean fixation rate observed in 100,000,000 simulated populations of size *N* = *Nf* = 1,000 under no viability selection (*Ns* = 0, brown), weak viability selection (*Ns* = 1, blue), and strong viability selection (*Ns* = 10, green). Dashed horizontal lines indicate the corresponding fixation probabilities of the underdominant allele under random mating. Fixation probabilities under the two models of assortative mating are equal either when *n* = 1 and the selfing probability is zero, or when *n* → ∞ and the selfing probability equals one. Under classical assortative mating the fixation probability interpolates smoothly between these two limits. However, the *n*-choice model has a complex effect on fixation probabilities between these two limiting cases. At small *n* > 1, the fixation probability is vanishingly small, and it depends on the strength of viability selection; whereas at large *n* the fixation probability depends only on *n* and it can even exceed the fixation probability under random mating. Thus *n*-choice assortative mating can either impede or facilitate fixation of an underdominant allele.

Nevertheless, as the number of mating opportunities increases further we find a surprising result: viability selection against heterozygotes and *n*-choice assortative mating — two selective forces that each act against rare alleles — interact paradoxically to increase the fixation probability of new mutant alleles. At sufficiently large, finite *n*, the fixation probability of a new underdominant allele under *n*-choice mating exceeds even that under random mating. For example, when selection is strong as *Ns* = 10, *n*-choice mating is beneficial to the rare underdominant allele provided the number of mating opportunities is a substantial fraction of the population, e.g. *n* = 0.2*N*. The stronger viability selection acts against heterozygotes, the smaller *n* is required for mate choice to facilitate fixation, relative to random mating.

This counterintuitive interaction between viability and sexual selection occurs because the manifold of equilibrium heterozygosity is reduced to 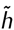(*p*) ≈ 0, for *n* large. Thus, there are few or no heterozygotes in the population on which viability selection can act. When there are many mating opportunities, a heterozygote parent is likely to find another heterozygote to mate with, and through Mendelian segregation this mating results in half the frequency of heterozygous progeny in each successive generation: roughly half of the minor alleles present in heterozygotes are transferred to homozygotes at each mating. Thus with a sufficient number of mating opportunities, the mating system effectively hides hybrids from the eyes of underdominant selection.

## 5. DISCUSSION

We have studied a mechanistic, one-parameter model of assortative mating that naturally spans the two classical extremes of random mating and complete assortment. The *n*-choice model fulfills the realistic condition that individuals can survey only a limited number of prospective mates. This simple formulation of mate selection nonetheless induces a complex form of sexual selection against rare alleles. In some regimes the induced selection is strong enough to virtually prevent the fixation of rare alleles. If the locus guiding mate choice is also subject to underdominant viability selection then, provided the number of mating opportunities is large, the *n*-choice mating system can mask the effects of viability selection, greatly elevating the fixation rate of underdominant alleles in comparison to random mating.

The *n*-choice mating model provides a qualitatively different resolution to the puzzle of the observed fixation of underdominant alleles between populations. In a well-mixed population the fixation rate of an underdominant mutation decreases rapidly with population size. Wright’s island model resolves this puzzle by exogenously subdividing the population into demes, so that fixation depends on the size of the demes rather than the whole population. For underdominant loci that also inflence mate choice, the *n*-choice model can effectively decouple the fixation rate from population size without imposing a fixed population structure. Although both classical models of structured populations or partial selfing and the *n*-choice model facilitate fixation of underdominant alleles by suppressing heterozygosity, the mechanism and consequences of mating structure differ. In structured populations or partial selfing the inbreeding coefficient is exogenously fixed and it does not depend on allele frequency, whereas in the *n*-choice model the mating structure depends on the frequency of the rare allele so that, in particular, mating still occurs at random in monomorphic populations.

The precipitous drop in fixation probability of a novel mutant between random mating and *n*-choice mating, from *n* = 1 to *n* = 2 mating opportunities, is surprising. From the gestating parent’s perspective, mate choice can only help rare alleles, as carriers of a rare allele copy have a greater chance of finding their own genotype to mate with and thus a lower chance of heterozygous offspring. However this gain of female function comes at a cost to male function. Because common types nearly always find their mates, but rare genotypes are more likely to settle for a common type, rare males are selected against. The relative strengths of reduced male fitness, heterozygote viability selection, and increased female fitness shift as the number of mating opportunities *n* increases, making preferred mates more easily accessible. Female fitness is enhanced through reduction of viability selection against their offspring, and male fitness is less affected with higher values of *n*. The exact crossover point where fixation becomes more likely under *n*-choice mating than random mating depends on the strength of viability selection against heterozygotes, *Ns*, and the rate of participation in the mating system, *Nf*. Fixation probabilities that depend non-monotonically on a physical parameter are unusual, but they have also arisen, for different reasons, in models of subdivided populations with extinction and recolonization[Michalakis and Olivieri, 1993; Roze and Rousset, 2003].

The *n*-choice model can be seen as a (degenerate) case of the ‘best-of-*n*′ model introduced to study the efficiency of mate choice mechanisms[Janetos, 1980]. Best-of-*n* models have been used in deterministic and stochastic settings to study the maintenance and efficiency of sexual selection[Seger, 1985; Pomiankowski, 1987] and speciation[Higashi *et al.*, 1999; Arnegard and Kondrashov, 2004; Servedio and Bürger 2014]. Incorporating best-of-*n* mating into population-genetic models is known to produce different conclusions than under fixed relative preference assortative mating[Seger, 1985; Kuijper *et al.*, 2012] for the efficiency of speciation[Arnegard and Kondrashov, 2004; M’Gonigle and FitzJohn, 2010; Servedio and Bürger 2014]. Despite these findings, most work on the fate of alleles in finite populations neglects mechanisms of non-random mate choice. The methods we have used to study the effects of *n*-choice mating in finite populations may be extended to other frequency-dependent mechanisms of mate choice and to other forms of viability selection and dominance on alleles.

We have described the *n*-choice model in the context of diploid parents, but minor variants of the model show similar behavior. For example, an analogous model in which a diploid parent censuses haploid microgametes, as occurs in flowering plants, has a different fixation probability in the limit *n* → ∞ (see Supplementary Fig. S1), but the same qualitative behavior remains: *n*-choice mating still induces sexual selection against rare types that can interact with underdominant viability selection to either impede or facilitate fixation of new mutants. Alternatively, if we prohibit selfing in the diploid model, *n*-choice mating is still more effective than random mating at fixing underdominant alleles at high *n* and *Ns* (Supplementary Fig. S2).

The physical interpretation of the pool of prospective mates in the *n*-choice model may differ between species, depending upon life history. In some species, *n* may count the number of possible matings over a lifetime (e.g., for semelparous species) or during a gestation period. One Borage species, for example, *Cryptantha flava* contains four ovules per flower, and the plant typically grows only one into a seed even when all are fertilized[Casper, 1984]. If some pairs of alleles have an embryonic lethal phenotype as hybrids, we can think of each flower as censusing four possible mates, testing each mating product for homozygosity and raising only homozygotes Thus the mating system of *C. flava* corresponds to the *n*-choice model with *n* = 4.

We introduced the parameter *f*, describing the proportion of the population that undergoes mate choice as opposed to clonal reproduction. Although this parameter was introduced for technical reasons, in order to produce a well-defined diffusion limit, even in finite models *Nf* has a natural, physical interpretation as the rate of mating: the average number of matings per generation, or the relative strength in altering gene frequencies by the mating system versus by genetic drift. One might naively assume that *f* is always unity in natural populations, and yet many plants such as some grasses and aspen reproduce sexually on a background of clonal reproduction. Genetic drift due to accidents of sampling can be interpreted at many levels, including sampling induced by the outcomes of mating; or stochastically induced by persistence to the next generation through longevity.

The complex sexual selection induced by the *n*-choice mating model and its counterintuitive interaction with underdominant viability selection remind us that relaxing population-genetic assumptions can radically alter allele frequency dynamics in surprising ways. The astounding diversity of life-histories across taxa provides ample motivation to rethink conclusions drawn from standard models of randomly-mating diploid populations.

## 6. MATERIALS AND METHODS

To derive a diffusion approximation we start with the standard Kolmogorov forward equation in two dimensions[Gardiner, 2009],

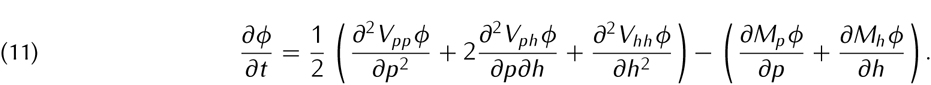

Here, *M*_*i*_ and *V*_*ij*_ represent the instantaneous mean and variance of allele frequency change. Assuming only a fraction *f* of the population undergo mating according to Eq. 4, the expected frequencies in the next generation, *p*′ and *h*′, are given by replacing *π* and *η* with *fπ* + (1 − *f*)*p* and *fη* + (1 − *f*)*h* in Eq. 5:

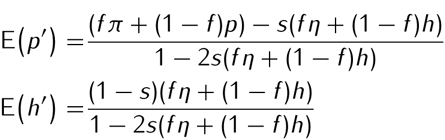

The mean change per generation is then *M*_*p*_ = E(*p*′) — *p* and *M*_*h*_, = *E*(*h*′) — *h*, which are rational functions in *p* and *h* of height 2*n*. We take the limit *N* → ∞, while holding both *N* = *ζ* and *Ns* = *γ* constant, so that *f* and *s* are small parameters. Thus only first-order terms survive in the Taylor expansion of *M*_*p*_ and *M*_*h*_, around (*f, s*) = (0, 0). We are left with

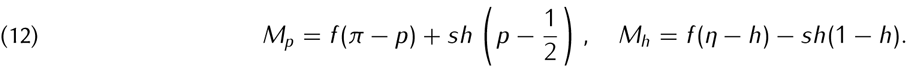

As *f* and *s* approach zero in this limit, so does the mean change in allele frequency, and E(*p*′) → *p* and E(*h*′) → *h*. Thus the variance-covariance matrix of allele frequency change approaches the multinomial variance-covariance matrix of sampling from the current allele frequencies. Thus the *V*_*ij*_ are given simply by Eq. 6: *V*_*pp*_(*p, h*) = Var(*p*), *V*_*hh*_(*p, h*) = Var(*h*), and *V*_*ph*_(*p, h*) = Cov(*p, h*), where *E*(*p*) = *p*. We rescale time in Eq. 11, taking *τ* = *t/N*:

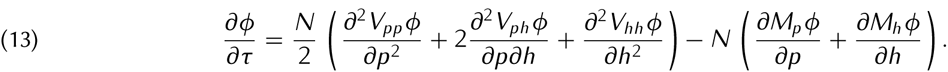

Writing Eq. 13 in terms of *π, p, η* and *h* and combining factors *Nf* = *ζ* and *Ns* = *γ* gives Eq. 7.

We compute the integral in Eq. 10 and the manifold of equilibrium heterozygosity (Eq. 8) numerically using Mathematica (Wolfram Research, Inc, Mathematica, Version 10.0.2.0 (2015), Champaign, IL, USA). We also wrote software in OCaml using the GNU Scientific Library to estimate fixation probabilities of the discrete model by explicit Monte Carlo simulation. The software is open source and available on GitHub (https://github.com/mnewberry/XXXXXXX).

## 7. ACKNOWLEDGEMENTS

We thank Brenda Casper, Erol Akçay, Tim Linksvayer, and Paul Sniegowski for important comments. We acknowledge funding from the Burroughs Wellcome Fund, the David and Lucile Packard Foundation, the James S. McDonnell Foundation, the Alfred P. Sloan Foundation, the U.S. Department of the Interior, the U.S. Army Research Office, and the National Institutes of Health.

---

**FIGURE S1.**
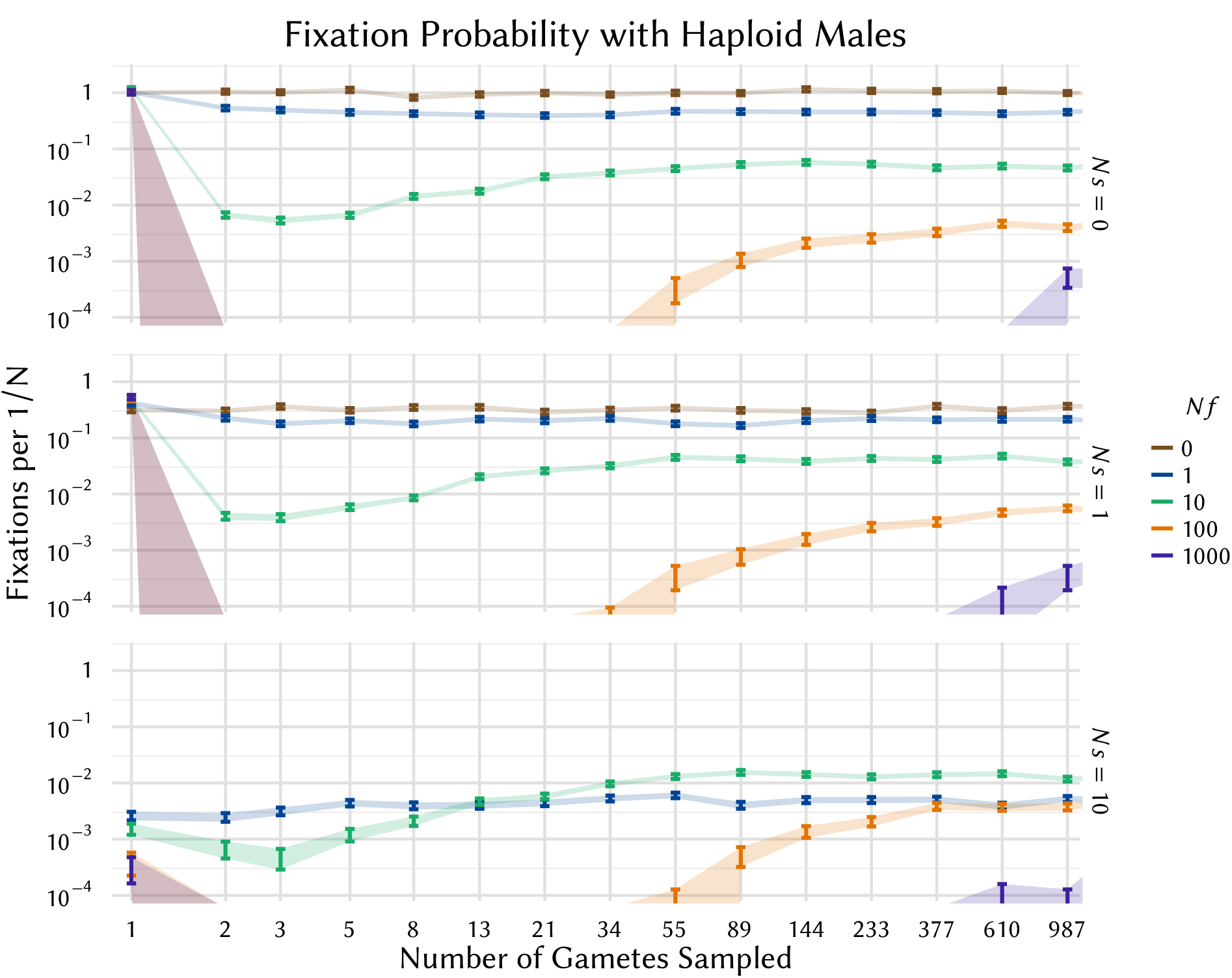
The fixation probability of one initial heterozygote in a model where females sample microgametes (sperm) attempting to raise homozygous offspring. This model is analogous to the *n*-choice model we study, but males are replaced by haplotypes. Females sample a limited number of gametes (*n*) and choose the first one that allows them to produce a homozygote, or, failing that, produce the heterozygote. The plot shows the fixation probability at different levels of viability selection against heterozygotes (*Ns*), and different rates of female participation in the mating system (*Nf*) on a background of clonal reproduction. Bands and error bars indicate the 95% confidence interval on the mean fixation rate in simulations with up to 100,000,000 runs in populations of size *N* = 1,000. For *Ns* = 10 and *Nf* = 0 (brown) the probabilities are below the range depicted. When females sample only one gamete (*n* = 1), the fixation probability is still roughly approximated by Eq. 1. At intermediate *n*, participation in the mating system induces strong fecundity selection against rare alleles. At large *n*, the fixation probability does not approach the neutral rate, because in order to form an initial mutant homozygote, an initial heterozygote must be chosen to reproduce, and it must also chose a mutant sperm instead of the more abundant wild-type. This depresses the fixation probability in the high-*n* limit relative to neutrality. Nonetheless, at large *n* and *Ns* the mating system can facilitate underdominant fixation.

**FIGURE S2.**
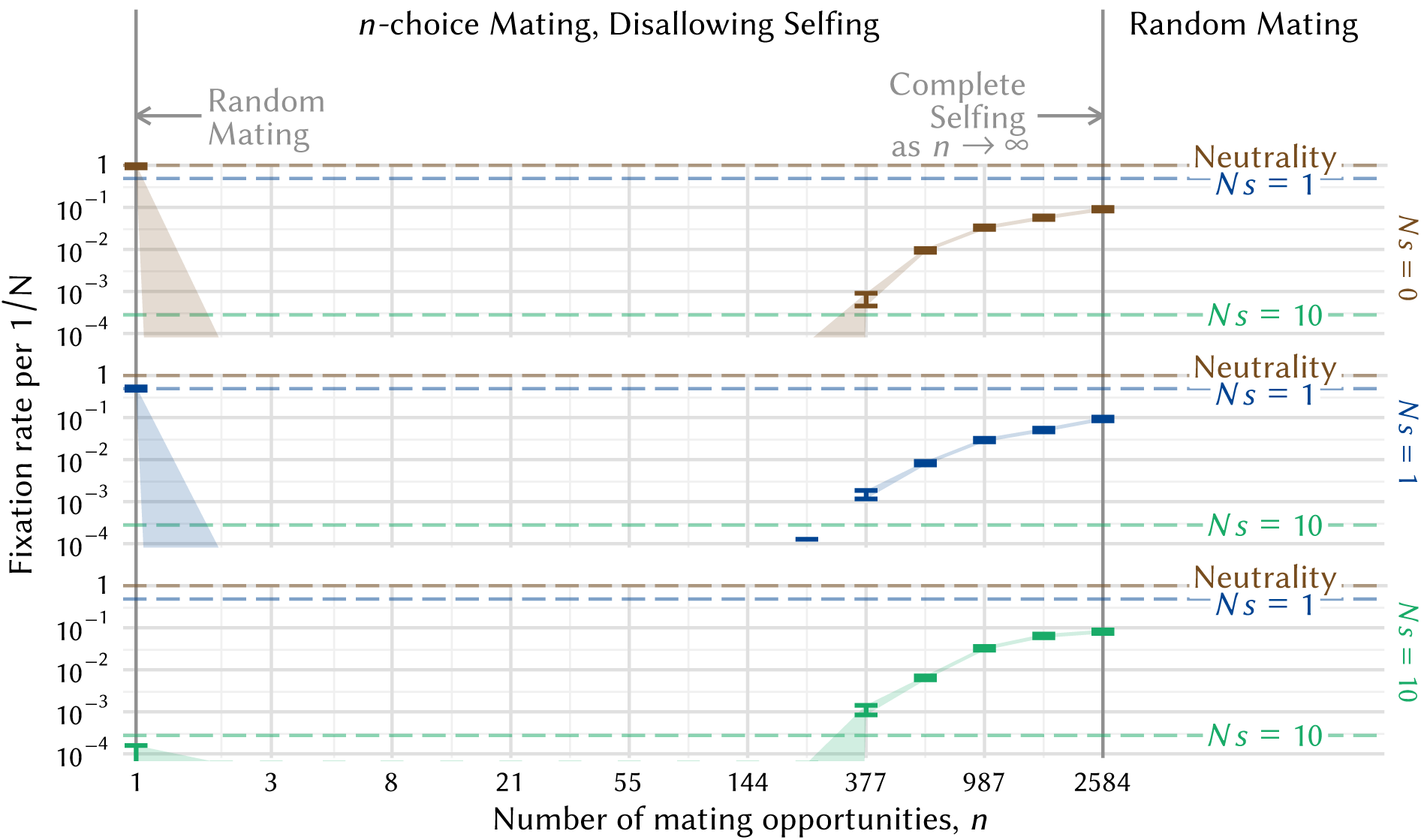
The effect of *n*-choice assortative mating on the fixation probability of an underdominant allele when self-mating is disallowed. The model and parameters are the same as those used in Figure 3, except that zygotes are drawn from a modified version of Eq. 2 which accounts for prohibition on self-mating. Vertical bars indicate the 95% confidence interval on the mean fixation rate observed in 100,000,000 replicate simulated populations of size *N = Nf* = 1,000 under no viability selection (*Ns* = 0, brown), weak viability selection (*Ns* = 1, blue), and strong viability selection (*Ns* =10, green). Dashed horizontal lines indicate the corresponding fixation probabilities of the underdominant allele under random mating. The asymptotic fixation probabilities at high *n* are depressed relative to neutrality because an initial homozygote must first drift to copy number higher than 1 before its own genotype is available for mating.

## REFERENCES

Philipp M Altrock, Arne Traulsen, and Floyd A Reed. Stability properties of underdominance in finite subdivided populations. PLoS Comput Biol, 7(11):e1002260–e1002260, 2011.

C Anderson, SC Wong, A Fuller, K Zigelsky, and RL Earley. Carotenoid-based coloration is associated with predation risk, competition, and breeding status in female convict cichlids (amatitlania siquia) under field conditions. Environmental Biology of Fishes, 98(4):1005–1013, 2015.

Malte Andersson and Leigh W Simmons. Sexual selection and mate choice. Trends in Ecology & Evolution, 21(6):296–302, 2006.

Malte B Andersson. Sexual selection. Princeton University Press, 1994.

Carlos F Arias, Astrid G Munoz, Chris D Jiggins, Jesus Mavarez, Eldredge Bermingham, and Mauricio Linares. A hybrid zone provides evidence for incipient ecological speciation in heliconius butterflies. Molecular ecology, 17(21):4699–4712, 2008.

Matthew E Arnegard and Alexey S Kondrashov. Sympatric speciation by sexual selection alone is unlikely. Evolution, 58(2):222–237, 2004.

NH Barton and S Rouhani. The probability of fixation of a new karyotype in a continuous population. Evolution, pages 499–517, 1991.

Jonathan D Blount, Neil B Metcalfe, Tim R Birkhead, and Peter FSurai. Carotenoid modulation of immune function and sexual attractiveness in zebra finches. Science, 300(5616):125–127, 2003.

Jonathan D Blount. Carotenoids and life-history evolution in animals. Archives of Biochemistry and Biophysics, 430(1):10–15, 2004.

Armando Caballero and William G Hill. Effects of partial inbreeding on fixation rates and variation of mutant genes. Genetics, 131(2):493–507, 1992.

Brenda B Casper. On the evolution of embryo abortion in the herbaceous perennial cryptantha flava. Evolution, pages 1337–1349, 1984.

Brian Charlesworth. Evolutionary rates in partially self-fertilizing species. American Naturalist, pages 126–148, 1992.

Tim Clutton-Brock and Katherine McAuliffe. Female mate choice in mammals. The Quarterly Review of Biology, 84(1):3–27, 2009.

Bernard J Crespi. Causes of assortative mating in arthropods. Animal Behaviour, 38(6):980–1000, 1989.

CF Curtis. Possible use of translocations to fix desirable genes in insect pest populations. Nature, 218:368–369, 1968.

Christian Damgaard. Fixation of advantageous alleles in partially self-fertilizing populations: the effect of different selection modes. Genetics, 154(2):813–821, 2000.

Charles L Epstein and Rafe Mazzeo. Degenerate diffusion operators arising in population biology. Princeton University Press, 2013.

Rui Faria and Arcadi Navarro. Chromosomal speciation revisited: rearranging theory with pieces of evidence. Trends in ecology & evolution, 25(11):660–669, 2010.

Crispin Gardiner. Stochastic Methods: A Handbook for the Natural and Social Sciences Springer Series in Synergetics. Springer, 2009.

Sergey Gavrilets. Fitness landscapes and the origin of species (MPB-41). Princeton University Press, 2004.

Sylvain Glémin. Extinction and fixation times with dominance and inbreeding. Theoretical population biology, 81(4):310–316, 2012.

Carol Goodwillie, Susan Kalisz, and Christopher G Eckert. The evolutionary enigma of mixed mating systems in plants: occurrence, theoretical explanations, and empirical evidence. Annual Review of Ecology Evolution, and Systematics, pages 47–79, 2005.

BS Haldane. A mathematical theory of natural and artificial selection. part ii the influence of partial self-fertilisation, inbreeding, assortative mating, and selective fertilisation on the composition of mendelian populations, and on natural selection. Biological Reviews, 1(3):158–163, 1924.

Philip W Hedrick. The establishment of chromosomal variants. Evolution, pages 322–332, 1981.

M Higashi, G Takimoto, and N Yamamura. Sympatric speciation by sexual selection. Nature, 402(6761):523–526, 1999.

Anthony C Janetos. Strategies of female mate choice: a theoretical analysis. Behavioral Ecology and Sociobiology, 7(2):107–112, 1980.

Chris D Jiggins, W Owen McMillan, Walter Neukirchen, and James Mallet. What can hybrid zones tell us about speciation? the case of heliconius erato and h. himera (lepidoptera: Nymphalidae). Biological Journal of the Linnean Society, 59(3):221–242, 1996.

Adam G Jones and Nicholas L Ratterman. Mate choice and sexual selection: what have we learned since darwin? Proceedings of the National Academy of Sciences, 106(Supplement 1):10001–10008, 2009.

Durrell D Kapan. Three-butterfly system provides a field test of müllerian mimicry. Nature, 409(6818):338–340, 2001.

Samuel Karlin. Comparisons of positive assortative mating and sexual selection models. Theoretical population biology, 14(2):281–312, 1978.

Jane P Kenney-Hunt, Ty T Vaughn, L Susan Pletscher, Andrea Peripato, Eric Routman, Kilinyaa Cothran, David Durand, Elizabeth Norgard, Christy Perel, and James M Cheverud. Quantitative trait loci for body size components in mice. Mammalian genome, 17(6):526–537, 2006.

Motoo Kimura. On the probability of fixation of mutant genes in a population. Genetics, 47(6):713, 1962.

Motoo Kimura. Diffusion models in population genetics. Journal of Applied Probability, 1(2):177–232, 1964.

Mark Kirkpatrick and Nick Barton. Chromosome inversions, local adaptation and speciation. Genetics, 173(1):419–434, 2006.

Mark Kirkpatrick. Sexual selection and the evolution of female choice. Evolution, pages 1–12, 1982.

Hanna Kokko, Michael D Jennions, and Robert Brooks. Unifying and testing models of sexual selection. Annual Review of Ecology Evolution, and Systematics, pages 43–66, 2006.

Bram Kuijper, Ido Pen, and FranzJ Weissing. A guide to sexual selection theory. Annual Review of Ecology, Evolution, and Systematics, 43:287–311, 2012.

Russell Lande and Douglas W Schemske. The evolution of self-fertilization and inbreeding depression in plants. i. genetic models. Evolution, pages 24–40, 1985.

Russell Lande. Effective deme sizes during long-term evolution estimated from rates of chromosome rearrangement. Evolution, 33:234–251, 1979.

Russell Lande. Models of speciation by sexual selection on polygenic traits. Proceedings of the National Academy of Sciences, 78(6):3721–3725, 1981.

Gary M Langham and C Benkman. Specialized avian predators repeatedly attack novel color morphs of heliconius butterflies. Evolution, 58(12):2783–2787, 2004.

James Mallet and Nicholas H Barton. Strong natural selection in a warning-color hybrid zone. Evolution, pages 421–431, 1989.

Leithen K M’Gonigle and Richard G FitzJohn. Assortative mating and spatial structure in hybrid zones. Evolution, 64(2):444–455, 2010.

Yannis Michalakis and Isabelle Olivieri. The influence of local extinctions on the probability of fixation of chromosomal rearrangements. Journal of evolutionary biology, 6(2):153–170, 1993.

Thomas Nagylaki. Introduction to theoretical population genetics, volume 142. Springer-Verlag Berlin, 1992.

Arcadi Navarro and Nick H Barton. Accumulating postzygotic isolation genes in parapatry: a new twist on chromosomal speciation. Evolution, 57(3):447–459, 2003.

Peter O’Donald. Genetic models of sexual selection, volume 44. Cambridge University Press Cambridge, 1980.

Daniel Ortíz-Barrientos, Brian A Counterman, and Mohamed AF Noor. Gene expression divergence and the origin of hybrid dysfunctions. Genetica, 129(1):71–81, 2007.

Sarah P Otto, Maria R Servedio, and Scott L Nuismer. Frequency-dependent selection and the evolution of assortative mating. Genetics, 179(4):2091–2112, 2008.

Todd L Parsons and Tim Rogers. Dimension reduction via timescale separation in stochastic dynamical systems. arXiv preprint arXiv: 1510.07031, 2015.

Andrew Pomiankowski. The costs of choice in sexual selection. Journal of theoretical Biology, 128(2):195–218, 1987.

Leslie Real. Search theory and mate choice. i. models of single-sex discrimination. American Naturalist, pages 376–405, 1990.

R Guy Reeves, Jarosław Bryk, Philipp M Altrock, Jai A Denton, and Floyd A Reed. First steps towards underdominant genetic transformation of insect populations. PLoS ONE, 9(5):e97557, 2014.

Loren H Rieseberg. Chromosomal rearrangements and speciation. Trends in Ecology & Evolution, 16(7):351–358, 2001.

Denis Roze and François Rousset. Selection and drift in subdivided populations: a straightforward method for deriving diffusion approximations and applications involving dominance, selfing and local extinctions. Genetics, 165(4):2153–2166, 2003.

Kristina M Sefc, Alexandria C Brown, and Ethan D Clotfelter. Carotenoid-based coloration in cichlid fishes. Comparative Biochemistry and Physiology Part A: Molecular & Integrative Physiology, 173:42–51, 2014.

Jon Seger. Unifying genetic models for the evolution of female choice. Evolution, pages 1185–1193, 1985.

Maria R Servedio and Reinhard Bürger. The counterintuitive role of sexual selection in species maintenance and speciation. Proceedings of the National Academy of Sciences, 111(22):8113–8118, 2014.

Steven P Sinkins and Fred Gould. Gene drive systems for insect disease vectors. Nature Reviews Genetics, 7(6):427–435, 2006.

Montgomery Slatkin. Fixation probabilities and fixation times in a subdivided population. Evolution, pages 477–488, 1981.

Lisa M Smith, Kirsten Bomblies, and Detlef Weigel. Complex evolutionary events at a tandem cluster of arabidopsis thaliana genes resulting in a single-locus genetic incompatibility. PLoS genetics, 7(7):e1002164, 2011.

Alexander J Stewart, Robert M Seymour, Andrew Pomiankowski, and Max Reuter. Under-dominance constrains the evolution of negative autoregulation in diploids. PLoS Comput Biol, 9(3):e1002992, 2013.

Michael C Whitlock. Fixation of new alleles and the extinction of small populations: drift load, beneficial alleles, and sexual selection. Evolution, 54(6):1855–1861, 2000.

Michael C Whitlock. Fixation probability and time in subdivided populations. Genetics, 164(2):767–779, 2003.

Daniel D Wiegmann, Steven M Seubert, and Gordon A Wade. Mate choice and optimal search behavior: fitness returns under the fixed sample and sequential search strategies. Journal of theoretical biology, 262(4):596–600, 2010.

Sewall Wright. Evolution in mendelian populations. Genetics, 16(2):97, 1931.

Sewall Wright. On the probability of fixation of reciprocal translocations. American Naturalist, pages 513–522, 1941.

Sewall Wright. The shifting balance theory and macroevolution. Annual review of genetics, 16(1):1–20, 1982.

